# INFLAMMATORY ACTIVATION, EPIGENETIC SIGNATURE AND MORPHO-STRUCTURAL ALTERATION IN A COHORT OF PATIENTS WITH CHRONIC HEART FAILURE AND PERMANENT ATRIAL FIBRILLATION: A CROSS-SECTIONAL STUDY

**DOI:** 10.1101/2025.04.29.651356

**Authors:** Mario Daidone, Carlo Domenico Maida, Gaetano Pacinella, Stefania Scaglione, Federica Todaro, Maria Grazia Puleo, Rosaria Maria Pipitone, Giulia Lupo, Luisa Agnello, Marcello Ciaccio, Giuseppe Armentaro, Angela Sciacqua, Tiziana Di Chiara, Domenico Di Raimondo, Stefania Grimaudo, Alessandra Casuccio, Antonino Tuttolomondo, MEDIREG (Mediterranean Research Group)

## Abstract

**Background:** Atrial fibrillation (AF) is a complex disorder involving inflammatory, morpho-structural, and genetic factors. Emerging evidence suggests inflammation plays a pivotal role in AF initiation, maintenance, and outcomes.

**Methods:** We conducted a cross-sectional case-control study to assess whether, among patients with chronic heart failure (CHF) and permanent AF, serum inflammatory cytokines, metabolic parameters, epigenetic factors, and echocardiographic markers of structural remodeling are interrelated. Eighty-two consecutive CHF patients with permanent AF and 82 CHF patients with sinus rhythm and no AF history were enrolled from the Internal Medicine and Stroke Care Unit at “P. Giaccone” Hospital, Palermo, between January 2020 and May 2022.

**Results:** AF patients were older and exhibited higher left atrial volume index (LAVI), reduced left atrial (LA) strain, increased relative wall thickness (RWT), and lower ejection fraction (EF%). Renal markers, including estimated glomerular filtration rate (eGFR) and microalbuminuria, were significantly different. Inflammatory markers such as C-reactive protein (CRP), interleukin 6 (IL-6), interleukin 8 (IL-8), and NT-proBNP were elevated in the AF group. Multivariable logistic regression identified LAVI, LA strain, microalbuminuria, albumin/creatinine ratio (ACr), and protein/creatinine ratio (PCr) as independent predictors of AF. Correlation analyses demonstrated strong associations between echocardiographic, inflammatory, and renal parameters. No significant differences in circulating miRNAs were observed.

**Conclusions:** These findings highlight the intricate interconnection between atrial remodeling, renal dysfunction, and systemic inflammation in AF pathogenesis. A better understanding of these mechanisms may inform future risk stratification strategies and support the development of more effective, targeted therapies in patients with CHF.

## BACKGROUND

Atrial fibrillation (AF) is the most frequent of the supraventricular arrhythmias, with a prevalence in the general population of up to 1%. This figure rises significantly with increasing age, reaching 8% in the elderly over 80 years old [1].

It is characterized by chaotic and disordered atrial electrical activity that causes dyssynchronous atrial contraction and equally irregular conduction of the electrical impulse to the ventricles. The prevailing hypothesis on the genesis of arrhythmia is that there are electrical triggers that propagate through reentry circuits located in vulnerable areas of atrial myocardium, inducing chaotic electrical and consequently mechanical activity (one of these myocardial sites is located close to the pulmonary veins) [2]

The importance of this condition, as well as its weight in terms of public health, derives from the impact that atrial fibrillation has on the morbidity and mortality of patients suffering from it: known consequences of this disease are embolic stroke and heart failure. In addition to age, many risk factors contribute to the onset of AF: arterial hypertension, obstructive sleep apnoea and hypopnoea syndrome (OSAHS), valvular defects, cardiomyopathies, and several others [3,4].

Knowledge and understanding of AF over the years have led to reconsideration of it, no longer as a pure cardiac disease but as a multisystem disease: recent and sensationalistic, in this sense, is the definition of AF as an “inflammatory disease” due to the finding that systemic inflammation plays a role in the onset, maintenance, and outcome of arrhythmia. Our group, in this sense, has documented how the “inflammatory milieu” represented by a large group of cytokines is fundamental in the development of the disease, and that the inflammatory substrate, contrary to previous thinking, is not associated with an underlying structural heart disease, but with the arrhythmia “per se” [5].

In addition to the risk factors and predisposing conditions already listed, it has been hypothesised over the last few decades that AF has a genetic basis, and genome-wide association studies (GWAS) have shown the existence of polymorphisms and mutations that explain the inheritance that characterises some forms of AF [6].

Obviously, talking about genetic components and inheritance, it does not mean that AF has a vertical transmission or that it follows the rules of Mendelian genetics, but it refers to genetic mutations and polymorphisms of genes making proteins that regulate various cellular activities, whose interaction with environmental factors and in the presence of favourable conditions can lead to the onset of the disease [7].

The evolution of this concept, therefore, has been epigenetics: the onset of the disease depends on genetic factors that are not sufficient to determine it and the interaction of these with the external environment is required: this interaction depends, in turn, on non-predictable factors such as the individual’s lifestyle, and this gives rise to a phenotype that does not mirror the genotype, even though it derives from it. The term ‘epigenetics’, in fact, identifies a change in gene expression that does not depend on a change in the gene itself, but on the way it is expressed [8].

Among the various mechanisms by which the cell regulates gene expression, in addition to DNA methylation and histone modification, the role of non-coding RNAs (miRNAs) is increasingly being investigated, although in atrial fibrillation and cardiovascular disease in general their role has not yet been fully elucidated.

miRNAs are non-coding RNA fragments of 19-25 nucleotides produced by RNA polymerase II from the DNA strand during the process of transcription. They are not then translated into proteins, but have a post-transcriptional regulatory role on messenger RNAs (mRNAs) by binding at specific binding sites on them (3’ untranslated regions - 3’UTRs) preventing mRNAs from being translated or promoting their degradation [9,10].

To date, the scientific literature has explored and detailed the role of selected miRNAs in atrial fibrillation: miR-30 and miR-133 down-regulate TGF-beta gene expression[11], while miR-29 down-regulates collagen and fibronectin production[12]. Their negative regulation leads to an accentuation of fibrotic processes and cardiac remodelling, which are fundamental for maintaining and perpetuating arrhythmia.

Furthermore, miR-26 shares the antifibrotic action with the previous miRNAs and also seems to have a further role in down-regulating inward-rectifier K(+) current (IK1), present in cardiomyocytes and fibroblasts and able to promote the development of congestive heart failure in patients with AF as it supports the maintenance of chaotic and irregular electrical activity [13].

MiR-1 is the most abundantly expressed miRNA at the level of atria and ventricles, its down-regulation has a pro-arrhythmic effect through modulation of IK1 by up-regulation of potassium inwardly rectifying channel subfamily J member 2 (KCNJ2) [14].

Up-regulation of miR-21 results in worsening of fibrosis in patients with AF: through phosphorylation of signal transducer and activator of transcription 3 (STAT 3); moreover, cardiac fibroblasts stimulated by interleukin-6 (IL-6) increase miR-21 expression and STAT 3 phosphorylation, again confirming the crucial role of inflammation and its impact on epigenetic regulation as well [15].

Because fibrosis is the outcome of a number of molecular mechanisms that depend on inflammation and epigenetic modulation documented in AF, structural cardiac alterations are a common consequence. There are many ways to detect these alterations, and among these certainly include “atrial strain”: it assesses myocardial deformability (via “speckle tracking” technique, which allows to study shifts of segments and points of the heart muscle and to calculate myocardial deformation), which is inversely proportional to the degree of fibrosis (measured by MRI or detected by biopsy examination) [16].

## AIMS OF THE STUDY

Based on this premises, our group aims to conduct a case-control cross-sectional study with the objective of assessing whether in a cohort of patients with CHF and permanent AF:

- There are documented associations between serum levels of inflammatory cytokines, metabolic variables and echocardiographic variables as indices of fibrotic/structural damage.
- upregulation or downregulation of specific Mi-RNAs, correlates with serum levels of inflammatory cytokines in order to suggest the existence of pathways that may link inflammation to atrial fibrosis present in patients with permanent AF.

## MATHERIALS AND METHODS

Between January 2020 and May 2022, we enrolled all 82 consecutive patients with CHF and permanent AF and 82 consecutive patients with CHF and the same risk factors but with sinus rhythm and no known history of previous AF, consecutively admitted to the Internal Medicine with Stroke Care Ward of the “P. Giaccone” Hospital of Palermo January 2020 to May 2022. The diagnosis of AF was made based on at least one 12-lead electrocardiogram and according to the current guidelines [17]. Patients with AF of >12 months in duration and ≥ 1 attempt at electrical cardioversion to restore normal sinus rhythm were considered to have permanent AF. Hypertension was defined according to the 2018 ESH/ESC guidelines [17]. Type 2 diabetes mellitus was determined using a clinically based algorithm that considered age at onset, presenting weight and symptoms, family history, onset of insulin treatment, and history of ketoacidosis[18]. Hypercholesterolemia was defined as total serum cholesterol ≥200mg/dL, and hypertriglyceridemia as total serum triglyceride ≥150mg/dL on the basis of the National Cholesterol Education Program–Adult Treatment Panel III reports [19]

Chronic heart failure was defined according to the current ESC guidelines [20]

As controls, among the consecutive subjects admitted to our ward between January 2020 and May 2022, we enrolled those with CHF and the same risk factors of AF subjects but with sinus rhythm and no known history of previous AF.

Patients with other subtypes of atrial fibrillation (paroxysmal, persistent, long-standing persistent) or at least one of the following exclusion criteria were excluded from the cohorts of case and controls: acute heart failure, active neoplastic disease, chronic inflammatory diseases, acute infectious diseases, severe renal insufficiency (eGFR estimated by CKD-EPI <30ml/min/m2), severe liver failure, chronic steroid/anti-inflammatory treatment.

For AF subjects and controls we evaluated medical history, 12-lead ECG, and two-dimensional and pulsed Doppler echocardiography at admission.

### Laboratory analysis

Nonfasting blood samples were obtained by venipuncture. The collected material was centrifuged at 1700 *g*/relative centrifugal force, after which citrate, EDTA-, heparin-, and Trasylol-plasma were separated, as well as blood serum. Buffy coats were collected from EDTA tubes to enable an analysis of genetic factors. Dimethylsulfoxide was added to an additional EDTA tube for cryopreservation of blood cells. All blood aliquots were subsequently stored at −80°C within 2 hours after venipuncture.

### Determination of IL-6 serum concentration

The serum IL-6 concentration was measured in samples using an enzyme-linked immunosorbent assay (Intertest 6; Genzyme, Boston, MA) according to the kit procedure. The limit of detection of the test was 76 pg/ml, and lower levels were considered undetectable.

### Determination of IL-8 serum concentration

The serum IL-8 concentration was measured in samples using an enzyme-linked immunosorbent assay (Invitrogen; ThermoFischer, Whaltam, MA) according to the kit procedure. The limit of detection of the test was 15.6 pg/ml, and lower levels were considered undetectable.

### Determination of TNF-a serum concentration

The serum TNF-a concentration was measured in samples using an enzyme-linked immunosorbent assay (Invitrogen; ThermoFischer, Whaltam, MA) according to the kit procedure. The limit of detection of the test was 15.6 pg/ml, and lower levels were considered undetectable.

### Determination of MCP-1 serum concentration

The serum MCP-1 concentration was measured in samples using an enzyme-linked immunosorbent assay (Invitrogen; ThermoFischer, Whaltam, MA) according to the kit procedure. The limit of detection of the test was 15.6 pg/ml, and lower levels were considered undetectable.

### Determination of serum CRP concentration

The CRP concentration was measured using a fluorescence polarization immunoenzymatic assay (Abbott Laboratories, Chicago, IL). The limit of detection of the CRP assay is 5 mg/litre.

### RNA isolation Polymerase chain reaction-based analysis

Total RNA was purified from serum sample of 30 patients using the commercially available miRNeasy Serum/ Plasma Kit (cat. Number 217204; Qiagen) and miRCURY LNA RNA Spike-in kit (cat. Number 339390; Qiagen) according to the manufacturer’s instructions.

For miRNA expression, 1 μg of RNA was retro-transcribed to cDNA using miRCURY LNA kit (cat. Number 339340; QIAGEN) according to manufacturer’s recommendations. Quantitative Real Time PCR was performed using miRCURY LNA SYBR Green PCR kit; cat. Number 339346; Qiagen) and the Custom_miRCURY PCR Panel (96-Well Format, Cat. Number 339330, YCA41201, Qiagen). The plates contained primers to analyze 3 targets, has-miR-1-3p, has-miR26a-5p and has-miR-21-5p; and 5 housekeeping genes: has-miR-93-5P, has-miR-191-5p, has-miR-425-5p, has-miR-451a and has-miR-23a-3p, as reported in Supplementary Table 1.

**Table 1:**
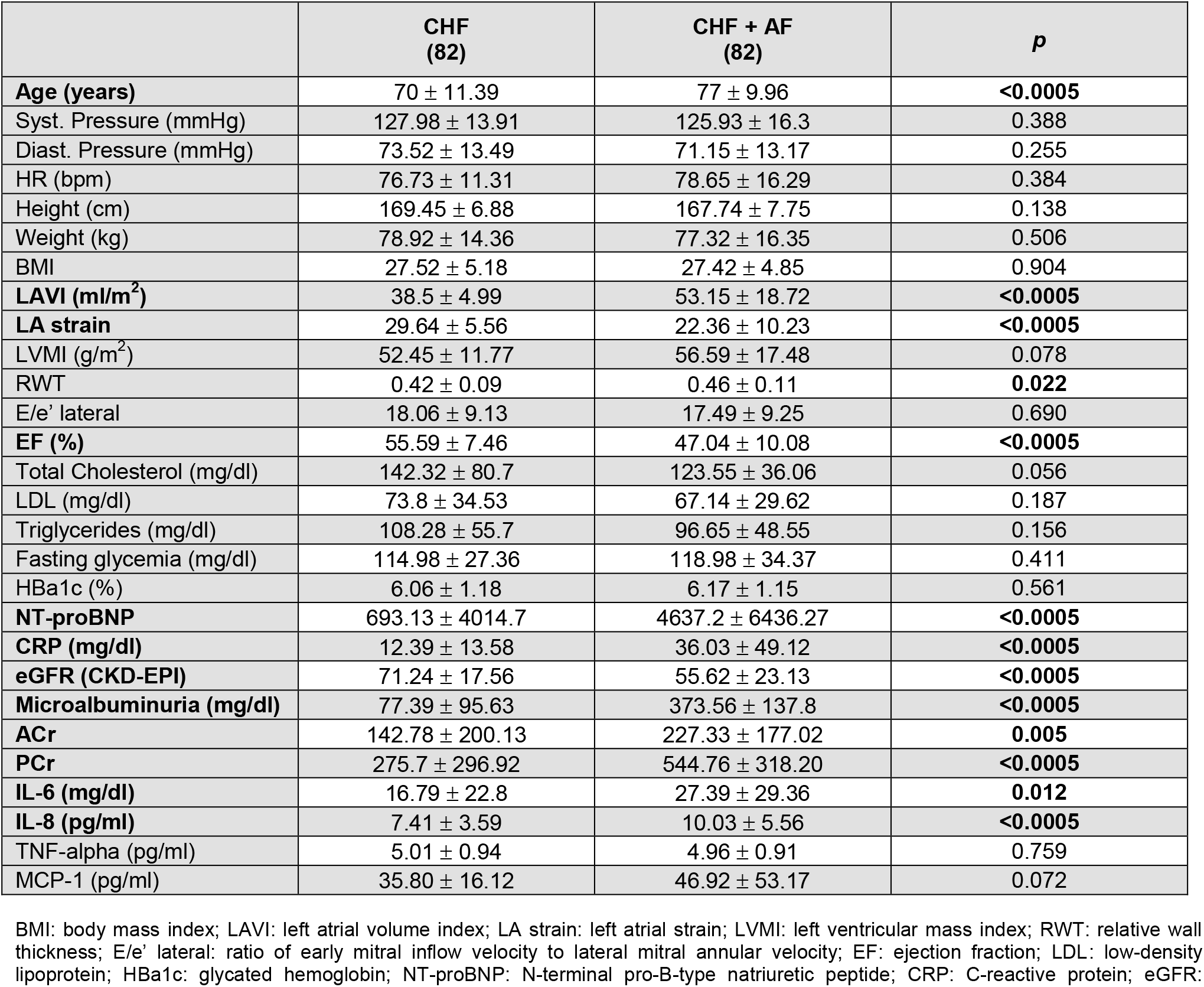
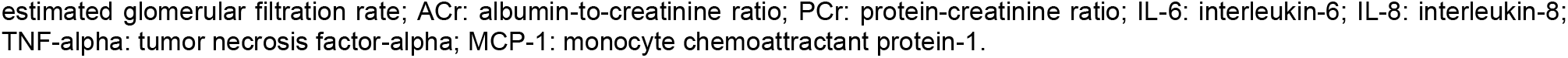
Descriptive analysis of anthropometric, clinical, echocardiographic and laboratory variables of patients with CHF and CHF + AF.

|  | CHF<br>(82) | CHF + AF<br>(82) | *p* |
| --- | --- | --- | --- |
| **Age (years)** | 70 ± 11.39 | 77 ± 9.96 | **<0.0005** |
| Syst. Pressure (mmHg) | 127.98 ± 13.91 | 125.93 ± 16.3 | 0.388 |
| Diast. Pressure (mmHg) | 73.52 ± 13.49 | 71.15 ± 13.17 | 0.255 |
| HR (bpm) | 76.73 ± 11.31 | 78.65 ± 16.29 | 0.384 |
| Height (cm) | 169.45 ± 6.88 | 167.74 ± 7.75 | 0.138 |
| Weight (kg) | 78.92 ± 14.36 | 77.32 ± 16.35 | 0.506 |
| BMI | 27.52 ± 5.18 | 27.42 ± 4.85 | 0.904 |
| **LAVI (ml/m<sup>2</sup>)** | 38.5 ± 4.99 | 53.15 ± 18.72 | **<0.0005** |
| **LA strain** | 29.64 ± 5.56 | 22.36 ± 10.23 | **<0.0005** |
| LVMI (g/m<sup>2</sup>) | 52.45 ± 11.77 | 56.59 ± 17.48 | 0.078 |
| RWT | 0.42 ± 0.09 | 0.46 ± 0.11 | **0.022** |
| E/e' lateral | 18.06 ± 9.13 | 17.49 ± 9.25 | 0.690 |
| **EF (%)** | 55.59 ± 7.46 | 47.04 ± 10.08 | **<0.0005** |
| Total Cholesterol (mg/dl) | 142.32 ± 80.7 | 123.55 ± 36.06 | 0.056 |
| LDL (mg/dl) | 73.8 ± 34.53 | 67.14 ± 29.62 | 0.187 |
| Triglycerides (mg/dl) | 108.28 ± 55.7 | 96.65 ± 48.55 | 0.156 |
| Fasting glycemia (mg/dl) | 114.98 ± 27.36 | 118.98 ± 34.37 | 0.411 |
| HbA1c (%) | 6.06 ± 1.18 | 6.17 ± 1.15 | 0.561 |
| **NT-proBNP** | 693.13 ± 4014.7 | 4637.2 ± 6436.27 | **<0.0005** |
| **CRP (mg/dl)** | 12.39 ± 13.58 | 36.03 ± 49.12 | **<0.0005** |
| **eGFR (CKD-EPI)** | 71.24 ± 17.56 | 55.62 ± 23.13 | **<0.0005** |
| **Microalbuminuria (mg/dl)** | 77.39 ± 95.63 | 373.56 ± 137.8 | **<0.0005** |
| **ACr** | 142.78 ± 200.13 | 227.33 ± 177.02 | **0.005** |
| **PCr** | 275.7 ± 296.92 | 544.76 ± 318.20 | **<0.0005** |
| **IL-6 (mg/dl)** | 16.79 ± 22.8 | 27.39 ± 29.36 | **0.012** |
| **IL-8 (pg/ml)** | 7.41 ± 3.59 | 10.03 ± 5.56 | **<0.0005** |
| TNF-alpha (pg/ml) | 5.01 ± 0.94 | 4.96 ± 0.91 | 0.759 |
| MCP-1 (pg/ml) | 35.80 ± 16.12 | 46.92 ± 53.17 | 0.072 |

BMI: body mass index; LAVI: left atrial volume index; LA strain: left atrial strain; LVMI: left ventricular mass index; RWT: relative wall thickness; E/e' lateral: ratio of early mitral inflow velocity to lateral mitral annular velocity; EF: ejection fraction; LDL: low-density lipoprotein; HbA1c: glycated hemoglobin; NT-proBNP: N-terminal pro-B-type natriuretic peptide; CRP: C-reactive protein; eGFR:
estimated glomerular filtration rate; ACr: albumin-to-creatinine ratio; PCr: protein-creatinine ratio; IL-6: interleukin-6; IL-8: interleukin-8; TNF-alpha: tumor necrosis factor-alpha; MCP-1: monocyte chemoattractant protein-1.

In addition, each plate contains primers to analyze 4 UniSp RNA spike-in: UniSp2, UniSp4 (RNA isolation controls), UniSp3 (interplate calibrator assay) and UniSp6 (cDNA Synthesis control) as reported in Supplementary Table 1.

The mean threshold cycle was used for the calculation of relative expression using the has-miR-451a as the reference gene.

Data were expressed as fold change using 2 ^− ΔΔCt^ method referred to subjects as control group. Differences among experimental groups were analyzed by Student t-test. The student’s t-test for independent experiments is performed for testing differences in fold expression of genes between the experimental groups; the Bonferroni correction for multiple hypothesis testing was applied to t-test and P-value, false-positive error probability, was used as primary criterion for selection of genes (P-value cutoff of 0.05 for statistical significance) and then the fold-change was considered as a measure of biological significance.

The cycle threshold (ct) values were submitted to the Web-based PCR Array Data Analysis software (https://geneglobe.qiagen.com/it/analyze).

### Statistical analysis

Statistical analysis of quantitative and qualitative data, including descriptive statistics, was performed for all items. Continuous data are expressed as mean ±SD, unless otherwise specified. Baseline differences between groups were assessed by the chi-square test or Fisher exact test, as needed for categorical variables, and by the independent Student t test for continuous parameters if the data were normally distributed. Logistic regression analysis examined the correlation between patient characteristics (independent variables), and patient groups (dependent variable) in a simple model. All variables with a p-value lower than 0.5 at univariate analysis were ran in a multivariate linear regression model and the data were adjusted for BMI, dyslipidemia, diabetes, ACE inhibitors or ARB, Beta blockade, SBP and DBP.

## RESULTS

We enrolled a total of 82 subjects with permanent AF and CHF and 82 subjects with CHF in sinus rhythm without any history of AF. General and laboratory characteristics of patients with AF and of subjects in sinus rhythm and without any history of AF are listed in Table 1.

Subject with AF, compared to control, showed a significantly higher age (77.8 vs 69.6, p <0.0005), no differences have been shown for blood pressure, lipids levels, glycemic variables and anthropometric variables. With regard to echocardiographic structural variables, patients with atrial fibrillation showed higher mean values of LAVI (53.152±18,7 vs 38.5±4.99, p <0.0005), lower LA strain (22.357±10.23 vs 29.64±5.56, p<0.0005), higher RWT

(0.0465±0.11 vs 0.42±0.09, p 0.022), lower EF% (47.04±10.08 vs 55.59±7.46, p <0.0005). No difference was highlighted with regard to echocardiographic parameters indicative of left ventricular filling pressures (E/e’ lateral 18±9,13 vs 17,5±9,2, p= 0,69) Inherent to renal function variables AF patients showed a lower estimated glomerular filtration rate (CKD-EPI) (55.63±23.13 vs 71.23±17.56 ml/min, p<0.0005), a higher quantitative microalbuminuria (173.55±137.8 vs 77.38±95.63 mg/dl, p<0.0005), a higher albumin/creatinine ratio (ACr) (142.78±200.13 vs 125.47±127.7, p 0.005), a higher protein/creatinine ratio (PCr) (544.76±318.21 vs 275.69±296.92, 296.92, p<0.0005). Regarding serological values of inflammatory cytokines and congestion indexes, AF patients showed higher mean values of CRP (36.03±49.12 vs 12.38±13.59 mg/dl, p<0.0005), IL6 (27.39±29.36 vs 16.79±22.80 mg/dl, p 0.012), IL8 (10.03±5.56 vs 7.41±3.59 mg/dl, p<0.0005) and NTproBNP (4637.19±6436.27 vs 693.130±4014.70 mg/dl, p<0.0005). Interestingly, no significant differences were found in miRNA levels between AF patients and controls.

A Multiple logistic regression model, including all the significant baseline variables (P<0.05) predictive of AF, showed that LAVI (OR: 1.06; 95%CI: 1.004-1.119; p=0.035), LA strain (OR: 0.868; 95%CI: 0.786-0.958; p=0.005), microalbuminuria (OR: 1.008; 95%CI: 1.002-1.014; p=0.012), ACr (OR: 0.988; 95%CI: 0.979-0.997; p=0.006) and PCr (OR: 1.006; 95%CI: 1.001-1.011; p=0.009) were independently associated with the arrhythmia (table 2).

**Table 2:** Multiple logistic regression model including all the significant baseline variables (P<0.05) predictive of AF.

| Group<sup>a</sup> | B | std.error | Wald | gl | Sign. | Exp (B) | 95% confidence interval for Exp(B) |  |
| --- | --- | --- | --- | --- | --- | --- | --- | --- |
|  |  |  |  |  |  |  | Lower limit | Upper limit |
| Intercept | -7,235 | 4,192 | 2,979 | 1 | ,084 |  |  |  |
| Age | ,066 | ,039 | 2,761 | 1 | ,097 | 1,068 | ,988 | 1,154 |
| **LAVI** | ,058 | ,028 | 4,440 | 1 | **,035** | **1,060** | **1,004** | **1,119** |
| **LAStrain** | -,142 | ,050 | 7,924 | 1 | **,005** | **,868** | **,786** | **,958** |
| RWT | 5,004 | 3,582 | 1,951 | 1 | ,162 | 148,993 | ,133 | 166842,299 |
| EF (%) | -,020 | ,049 | ,161 | 1 | ,688 | ,981 | ,891 | 1,079 |
| NT-PROBNP | ,000 | ,000 | ,000 | 1 | ,994 | 1,000 | 1,000 | 1,000 |
| PCR | ,031 | ,019 | 2,514 | 1 | ,113 | 1,031 | ,993 | 1,071 |
| eGFR (CDKEPI) | -,032 | ,017 | 3,447 | 1 | ,063 | ,969 | ,937 | 1,002 |
| **Microalbuminuria (quant)** | ,008 | ,003 | 6,286 | 1 | **,012** | **1,008** | **1,002** | **1,014** |
| **Acr** | -,012 | ,004 | 7,478 | 1 | **,006** | **,988** | **,979** | **,997** |
| **PCr_A** | ,006 | ,002 | 6,759 | 1 | **,009** | **1,006** | **1,001** | **1,011** |
| IL6 | ,003 | ,014 | ,033 | 1 | ,857 | 1,003 | ,976 | 1,030 |
| IL8 | ,089 | ,075 | 1,407 | 1 | ,236 | 1,093 | ,944 | 1,266 |
| **Cardiological disease** | ,917 | ,402 | 5,196 | 1 | **,023** | **2,501** | **1,137** | **5,501** |
| Hypocinesia | 1,134 | ,831 | 1,861 | 1 | ,173 | 3,108 | ,609 | 15,848 |
| Arterial Hypertension | ,675 | ,900 | ,563 | 1 | ,453 | 1,965 | ,337 | 11,468 |
| a. Reference category is: 0. |  |  |  |  |  |  |  |  |

At Pearson analysis, in the subgroup of patients with AF we showed a positive correlation between LAVI and LVMI (r: 0.223; p=0.044), E/e’ (r: 0.434; p=0.000), segmental hypokinesia (r: 0.352; p=0.001) and a negative correlation with EF% (r: -0.304; p=0.006), a positive correlation between LA strain and segmental hypokinesia (r: 0.453; p=0.000) and a negative correlation with NTproBNP (r: -0.247 ; p=0.025), a positive correlation between LVMI and E/e’ (r: 0.300; p=0.006) and segmental hypokinesia (r: 0.384 ; p=0.000) and a negative correlation with EF% (r: -0.355; p=0.001), a positive correlation between E/e’ and segmental hypokinesia (r: 0.351; p=0.001) and NTproBNP (r: 0.306; p=0.005) and a negative correlation with EF% (r: -0.367; p=0.001). For what concerning serum cytokines we showed a positive correlation between MCP-1 levels and EF%. (r: 0.230; p=0.042) (table 3).

**Table 3:** Pearson analysis of correlation between echocardiographic findings and serum cytokines in the subgroup of patients with CHF and AF.

|  |  | LAVI | LA Strain | LVMI | RWT | EE | FE (%) | HYPOCINESIA | NT-PROBNP | IL6 | IL8 | TNF-a | MCP1 |
| --- | --- | --- | --- | --- | --- | --- | --- | --- | --- | --- | --- | --- | --- |
| **LAVI** | Pearson correlation | 1 | ,124 | **,223\*** | ,169 | **,434\*\*** | **-,304\*\*** | **,352\*\*** | ,127 | -,047 | -,019 | -,101 | -,111 |
|  | Sign. (2 tailed) |  | ,269 | **,044** | ,129 | **,000** | **,006** | **,001** | ,255 | ,685 | ,864 | ,371 | ,331 |
|  | N | 82 | 82 | 82 | 82 | 82 | 82 | 82 | 82 | 78 | 81 | 80 | 79 |
| **LAStrain** | Pearson correlation | ,124 | 1 | -,062 | -,132 | ,001 | -,009 | **,453\*\*** | **-,247\*** | ,111 | ,012 | ,066 | ,002 |
|  | Sign. (2 tailed) | ,269 |  | ,581 | ,237 | ,990 | ,933 | **,000** | **,025** | ,335 | ,912 | ,558 | ,988 |
|  | N | 82 | 82 | 82 | 82 | 82 | 82 | 82 | 82 | 78 | 81 | 80 | 79 |
| **LVMI** | Pearson correlation | **,223\*** | -,062 | 1 | ,045 | **,300\*\*** | **,355\*\*** | **,384\*\*** | ,190 | -,038 | -,123 | ,153 | -,049 |
|  | Sign. (2 tailed) | **,044** | ,581 |  | ,685 | **,006** | **,001** | **,000** | ,088 | ,739 | ,274 | ,175 | ,667 |
|  | N | 82 | 82 | 82 | 82 | 82 | 82 | 82 | 82 | 78 | 81 | 80 | 79 |
| RWT | Pearson correlation | ,169 | -,132 | ,045 | 1 | ,161 | -,027 | ,025 | -,039 | -,160 | -,097 | -,003 | -,081 |
|  | Sign. (2 tailed) | ,129 | ,237 | ,685 |  | ,148 | ,811 | ,820 | ,727 | ,162 | ,388 | ,980 | ,476 |
|  | N | 82 | 82 | 82 | 82 | 82 | 82 | 82 | 82 | 78 | 81 | 80 | 79 |
| EE | Pearson correlation | **,434\*\*** | ,001 | **,300\*\*** | ,161 | 1 | **-,367\*\*** | **,351\*\*** | **,306\*\*** | ,022 | -,152 | ,060 | -,131 |
|  | Sign. (2 tailed) | **,000** | ,990 | **,006** | ,148 |  | **,001** | **,001** | **,005** | ,851 | ,175 | ,595 | ,250 |
|  | N | 82 | 82 | 82 | 82 | 82 | 82 | 82 | 82 | 78 | 81 | 80 | 79 |
| FEPERC | Pearson correlation | -,304** | -,009 | -,355** | -,027 | -,367** | 1 | **-,501\*\*** | -,208 | -,030 | ,017 | ,066 | **,230\*** |
|  | Sign. (2 tailed) | ,006 | ,933 | ,001 | ,811 | ,001 |  | **,000** | ,061 | ,797 | ,881 | ,559 | **,042** |
|  | N | 82 | 82 | 82 | 82 | 82 | 82 | 82 | 82 | 78 | 81 | 80 | 79 |
| HYPOCINESIA | Pearson correlation | ,352** | ,453** | ,384** | ,025 | ,351** | -,501** | 1 | ,110 | ,205 | -,008 | -,037 | -,133 |
|  | Sign. (2 tailed) | ,001 | ,000 | ,000 | ,820 | ,001 | ,000 |  | ,323 | ,072 | ,943 | ,742 | ,242 |
|  | N | 82 | 82 | 82 | 82 | 82 | 82 | 82 | 82 | 78 | 81 | 80 | 79 |
| NTPROBNP | Pearson correlation | ,127 | -,247* | ,190 | -,039 | ,306** | -,208 | ,110 | 1 | -,017 | -,105 | ,007 | ,061 |
|  | Sign. (2 tailed) | ,255 | ,025 | ,088 | ,727 | ,005 | ,061 | ,323 |  | ,882 | ,349 | ,952 | ,592 |
|  | N | 82 | 82 | 82 | 82 | 82 | 82 | 82 | 82 | 78 | 81 | 80 | 79 |
| IL6 | Pearson correlation | -,047 | ,111 | -,038 | -,160 | ,022 | -,030 | ,205 | -,017 | 1 | ,071 | ,201 | ,081 |
|  | Sign. (2 tailed) | ,685 | ,335 | ,739 | ,162 | ,851 | ,797 | ,072 | ,882 |  | ,534 | ,078 | ,482 |
|  | N | 78 | 78 | 78 | 78 | 78 | 78 | 78 | 78 | 78 | 78 | 78 | 77 |
| IL8 | Pearson correlation | -,019 | ,012 | -,123 | -,097 | -,152 | ,017 | -,008 | -,105 | ,071 | 1 | -,072 | ,055 |
|  | Sign. (2 tailed) | ,864 | ,912 | ,274 | ,388 | ,175 | ,881 | ,943 | ,349 | ,534 |  | ,527 | ,631 |
|  | N | 81 | 81 | 81 | 81 | 81 | 81 | 81 | 81 | 78 | 81 | 80 | 79 |
| TNFALFA | Pearson correlation | -,101 | ,066 | ,153 | -,003 | ,060 | ,066 | -,037 | ,007 | ,201 | -,072 | 1 | -,003 |
|  | Sign. (2 tailed) | ,371 | ,558 | ,175 | ,980 | ,595 | ,559 | ,742 | ,952 | ,078 | ,527 |  | ,976 |
|  | N | 80 | 80 | 80 | 80 | 80 | 80 | 80 | 80 | 78 | 80 | 80 | 79 |
| MCP1 | Pearson correlation | -,111 | ,002 | -,049 | -,081 | -,131 | ,230* | -,133 | ,061 | ,081 | ,055 | -,003 | 1 |
|  | Sign. (2 tailed) | ,331 | ,988 | ,667 | ,476 | ,250 | ,042 | ,242 | ,592 | ,482 | ,631 | ,976 |  |
|  | N | 79 | 79 | 79 | 79 | 79 | 79 | 79 | 79 | 77 | 79 | 79 | 79 |

\*. The correlation is significant at the 0.05 level (two-tailed).
\*\*.. The correlation is significant at the 0.01 level (two-tailed).

## DISCUSSION

The pathophysiology of atrial fibrillation, especially in recent years, has been explored in detail in order to understand, in parallel to the electrophysiological alterations underlying it, the biological, neurohormonal and molecular mechanisms on which there is considerable academic interest with a view to prevention and treatment of the pathology. In this regard, biomarkers of fibrosis and inflammation, given that AF is now considered an inflammatory and systemic disease, are the targets on which the scientific community’s interest is focused [21]. Our observational cross-sectional study aimed to investigate the relationship between atrial fibrillation, inflammation, metabolic dysfunction and epigenetic factor in a cohort of patient with CHF.

The observed higher age in AF patients aligns with existing literature highlighting age as a significant risk factor for AF [22]. Echocardiographic structural abnormalities, such as increased LAVI, reduced LA strain, higher RWT, and decreased EF%, are consistent with previous studies linking these parameters to AF [23]. Elevated levels of inflammatory markers (CRP, IL6, IL8) and NTproBNP in AF patients are in line with the established association between AF and inflammation, as well as the role of NTproBNP in cardiac dysfunction [24–26]

In our study, we observed statistically significant differences in serum levels of inflammatory cytokines such as IL-8, IL-6, and PCR between atrial fibrillation (AF) patients and controls [27]. However, our multinomial logistic regression analysis did not reveal significant associations between these cytokines and AF diagnosis. Possible explanations may include the influence of confounding variables, the dynamic nature of cytokine levels, or the existence of other unexplored inflammatory markers [28]. Previous research has highlighted the complexity of AF pathophysiology, involving factors beyond cytokines, such as genetic predisposition and structural heart changes [29]. Further studies are warranted to comprehensively understand the intricate mechanisms underlying AF.

The multiple logistic regression model’s identification of LAVI, LA strain, EF%, microalbuminuria, ACr, and PCr as independent predictors of AF is supported by the literature on the importance of these parameters in AF risk prediction [30,31].

The Pearson analysis revealing correlations between echocardiographic parameters and serum cytokines in the AF subgroup underscores the interplay between structural and inflammatory aspects in AF pathophysiology [32]

The absence of statistically significant differences in the expression of specific miRNAs (hsa-miR-1-3p, hsa-miR-26a-5p, hsa-miR-21-5p) between fibrillating patients and controls may be attributed to several factors. Firstly, miRNA expression is highly context-dependent and can be influenced by various external factors such as comorbidities, medications, and lifestyle, which might have obscured discernible patterns in our study [33,34]. Additionally, the intricate regulatory networks involving these miRNAs may contribute to their multifaceted roles in different physiological and pathological conditions[35]. Previous studies have reported the dynamic nature of miRNA expression and its susceptibility to fluctuations, underscoring the need for longitudinal assessments to capture temporal changes accurately [36]. Our findings align with the growing recognition of the complexity surrounding miRNA dysregulation in diverse disease states. Furthermore, our patient cohort consisted of subjects with chronic heart failure with preserved EF (HFpEF). As widely known, diastolic dysfunction occurs when there is a problem with myocardial relaxation and/or stiffness that translates into an increased left ventricular filling pressures. The results of our study highlighted the presence of an altered diastolic function in both study groups with E/e’ values at lateral TDI (Doppler tissue imaging) suggestive of elevated left atrial pressure. The results of our study suggest the absence, between cases and controls, of a statistically significant difference regarding the expression of specific pro-fibrotic miRNAs could be interpreted as an expression of the fact that both study groups were characterized by the presence of a remarkable myocardial stiffness related to the presence of chronic diastolic heart failure and in this context, the presence of arrhythmia would not seem to increase the fibrotic burden to myocardium. To the best of our knowledge, this is an interesting finding that further emphasizes the complexity of HFpEF which, in recent decades has evolved from mere LV diastolic dysfunction to a heterogeneous clinical syndrome, in which cardiac and vascular anomalies, cardiovascular risk factors and comorbidities coexist in various combinations.

Atrial fibrillation (AF) and chronic kidney disease (CKD) are closely interrelated conditions with an increasing prevalence, sharing common risk factors such as age, obesity, diabetes mellitus, hypertension, and ischemic heart disease. A bidirectional relationship exists between these diseases: AF predisposes individuals to CKD, while CKD itself increases the risk of AF. Several shared pathophysiological mechanisms have been identified over time, particularly systemic inflammation and activation of the renin-angiotensin-aldosterone system (RAAS). Elevated serum levels of inflammatory cytokines characterize the early stages of CKD and are also well-documented in AF. Supporting this, the use of anti-inflammatory agents such as glucocorticoids and statins has been shown to reduce AF incidence and slow renal function decline by mitigating the impairment of estimated glomerular filtration rate (eGFR) [37,38]. RAAS activation represents another crucial link between AF and CKD, as it promotes inflammation, oxidative stress, and fibrosis through increased serum concentrations of cytokines, cell adhesion molecules, and profibrotic growth factors. This cascade contributes to both atrial structural remodeling and renal dysfunction [39,40]. Notably, RAAS activation also induces the TGF-beta 1/Smad pathway, which enhances oxidative stress and fibrosis. Evidence from trials using pirfenidone, a TGF-beta 1 inhibitor, has demonstrated reduced fibrosis in cardiac, pulmonary, renal, and hepatic tissues [41–43]. Consequently, RAAS inhibition has been associated with a slower progression of renal impairment and reduced cardiac remodeling in AF patients [44].

Our study reinforces this pathophysiological connection by identifying key renal biomarkers— microalbuminuria, albumin-to-creatinine ratio (ACr), and protein-to-creatinine ratio (PCr)—as independent predictors of AF, as demonstrated through multiple logistic regression analysis. Microalbuminuria, a marker of endothelial dysfunction and oxidative stress, serves as an indicator of vascular damage in AF patients, providing further evidence of the common pathophysiological substrate linking AF and CKD. These findings highlight the crucial role of systemic inflammation and vascular injury, beyond traditional risk factors, in the interplay between the two conditions. Further research is warranted to explore targeted therapeutic strategies aimed at mitigating their reciprocal progression.

In addition to the classic markers of inflammation about which the scientific literature has now revealed many curiosities, our interest focused on a chemotactic cytokine, monocyte chemoattractant protein-1 (MCP-1) also known as CCL2, member of the C-C chemokine family, and a potent chemotactic factor for monocytes. This chemokine is upregulated in patients with AF, and plays a role in atherogenesis and thrombosis; it is involved in the production of several matrix metalloproteinases: in subjects with higher, genetically predetermined levels of MCP1, the risk of cardioembolic stroke is 14% higher for each standard deviation increase in chemokine levels [45]. Several studies have documented that MCP1 levels are higher in AF patients over the age of 65, while its serum levels are lower in young people with AF and in elderly controls, thus suggesting that this cytokine may play an active role in the genesis of fibrosis; confirming this finding, MCP-1-induced proteins (MCPIP - MCP1 family proteins) levels correlated with two known markers of fibrosis which are PIIINP (procollagen type III N-terminal peptide) and ICTP (type I collagen C-terminal telopeptide).

## LIMITATION

This study is limited by the variability in medication use between AF patients and controls. The cross-sectional analysis is a limitation as it does not allow conclusions to be drawn about cause-and-effect relationships, and the relatively small sample size may also limit conclusions.

## CONCLUSION

Atrial fibrillation represents a multifaceted pathology, in which there are complex interrelationships between inflammatory, morpho-structural and genetic variables. The study’s findings contribute to the growing body of evidence linking age, echocardiographic parameters, renal function, and inflammatory markers to AF, providing a comprehensive understanding of the multifaceted aspects associated with this complex arrhythmia. However, further studies are needed to better clarify causal relationship between the numerous pathophysiological variables that influence the course and prognosis of patients with atrial fibrillation.

